# Crystal structure of the membrane (M) protein from a SARS-COV-2-related coronavirus

**DOI:** 10.1101/2022.06.28.497981

**Authors:** Xiaodong Wang, Yuwei Yang, Ziyi Sun, Xiaoming Zhou

## Abstract

The membrane (M) protein is the most abundant structural protein of coronaviruses including SARS-COV-2 and plays a central role in virus assembly through its interaction with various partner proteins. However, mechanistic details about how M protein interacts with others remain elusive due to lack of high-resolution structures. Here, we present the first crystal structure of a coronavirus M protein from *Pipistrellus* bat coronavirus HKU5 (batCOV5-M), which is closely related to SARS-COV-2 M protein. Furthermore, an interaction analysis indicates that the carboxy-terminus of the batCOV5 nucleocapsid (N) protein mediates its interaction with batCOV5-M. Combined with a computational docking analysis an M-N interaction model is proposed, providing insight into the mechanism of M protein-mediated protein interactions.

## Main Text

The virion of coronaviruses including SARS-COV-2 contains four structural proteins: S (spike), M (membrane), E (envelope) and N (nucleocapsid) proteins^1-4^. Among them, M protein is the most abundant protein component and plays a central role in virus assembly mainly as a scaffolding center^1,5-7^. In coronaviruses, M protein is the only multi-spanning transmembrane structural protein and has been shown to interact with various partner proteins, including all viral structural proteins (S, M, E, N)^1,6,8-15^ and some host factors involved in the interferon signaling pathway such as MAVS^16-19^. However, how M protein interacts with its partners remains largely unknown due to lack of high-resolution structures of M protein^20-23^. In this study, we aim to determine the crystal structure of a coronavirus M protein.

To obtain a suitable M protein for structural study, we screened expression of M protein genes from eight betacoronavirus strains^1,24^, including SARS-COV-2 (SARS2), SARS-COV (SARS) and MERS-COV (MERS). Among them, the M protein from *Pipistrellus* bat coronavirus HKU5 (batCOV5-M) displayed the best expression and biochemical properties, and was selected for further structural study. The batCOV5-M protein is closely related to SARS2-M, SARS-M and MERS-M proteins, sharing 37% sequence identity and 68% sequence similarity among them (Supplementary Fig. 1).

After extensive crystallization trials and refinement, batCOV5-M crystallized in lipidic cubic phase (LCP) and the crystals diffracted to ∼3 Å resolution. However, molecular replacement^25^ using an AlphaFold^26^-predicted SARS2-M model (termed SARS2-M_AF_)^21^ failed to yield a valid solution, suggesting that SARS2-M_AF_ may vary significantly from the experimental M structure. Therefore, we sought to resolve the phase problem by experimental means, such as single-wavelength anomalous diffraction (SAD) using heavy atoms^27^. Since batCOV5-M crystals grew only in the LCP glass sandwich^28^, which makes soaking the crystals with heavy atoms impossible, we attempted to co-crystallize batCOV5-M with heavy atoms, but unfortunately unsuccessful. Our next strategy was to crystallize the cytosolic carboxy-terminal domain (CTD) of batCOV5-M (batCOV5-M_CTD_, Supplementary Fig. 1) and solve its structure, which may serve as a searching template for molecular replacement of the full-length batCOV5-M. In the end, by fusing the two halves of a split superfolder GFP^29,30^ to the two termini of batCOV5-M_CTD_, the resulting batCOV5-M_CTD_-GFP was crystallized by the vapor-diffusion sitting-drop method and its structure solved to 3.42 Å (Supplementary Fig. 2a) by molecular replacement using the GFP structure (PDB: 2B3Q) as a searching model. Interestingly, four batCOV5-M_CTD_-GFP molecules assemble into a tetramer in the structure with M_CTD_ domains swapping β strands with each other, which resembles a “butterfly” shape (Supplementary Fig. 2b). Three monomer conformations of M_CTD_ were then extracted from the tetramer: conformer 1 with the β sandwich split into three parts (Supplementary Fig. 2c), conformer 2 with two parts (Supplementary Fig. 2d), and conformer 3 as an assembled M_CTD_ (termed batCOV5-M_CTD-xtal_, Fig. 1a and Supplementary Fig. 2e). Though it is tempting to postulate that domain swapping of CTD may play a role in regulating the M-M protein interaction^1^, further investigation is required. Meanwhile, we tried to solve the full-length batCOV5-M structure by molecular replacement using the three CTD conformations as templates, and eventually succeeded with the batCOV5-M_CTD-xtal_ structure. The final batCOV5-M model was determined to 3.21 Å (termed batCOV5-M_xtal_, Fig. 1b) and contained residues 13-199 of batCOV5-M, and the amino-terminus (12 residues) and the carboxy-terminus (21 residues) were not modeled due to weak electron densities.

**Figure 1.**
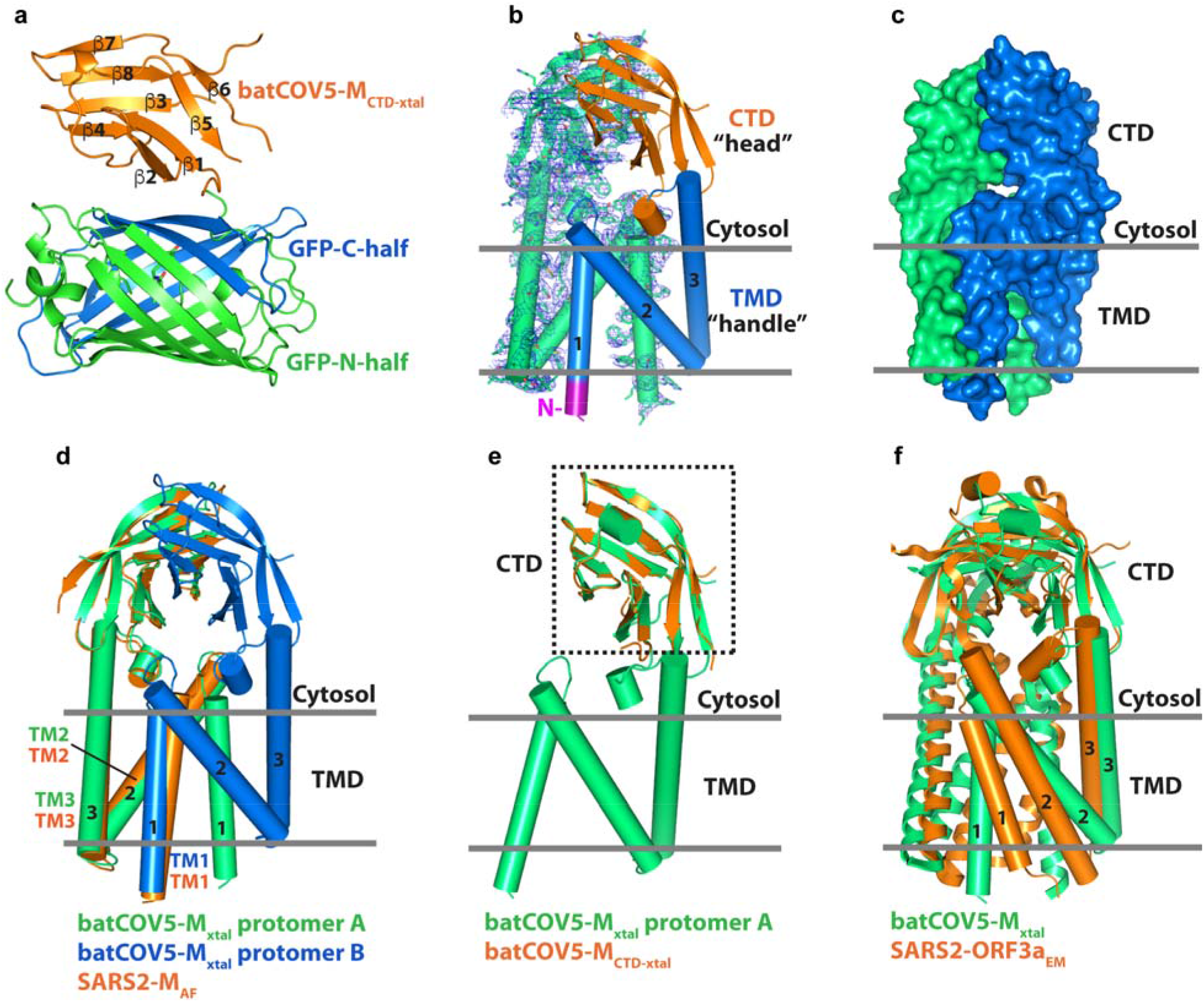
Crystal structure of M protein of the betacoronavirus batCOV5. (a) Crystal structure of batCOV5-M_CTD_ (in orange) fused with a split superfolder GFP (N-terminal half in green and C-terminal half in blue) rendered in the cartoon mode. The β strands of CTD are labeled from β1 to β8. (b) Crystal structure of batCOV5-M homodimer. For one protomer (in green), the 2F_o_-F_c_ map contoured at 1.2 σ level is shown as blue mesh. In the other protomer, the N-terminus, TMD and CTD are displayed in purple, blue and orange, respectively. The TM helices are labeled from 1 to 3 throughout this manuscript. The “head” and “handle” labels refer to an analogy to a pair of pincers in the main text. Relative membrane position is indicated by two grey lines throughout this manuscript. (c) The batCOV5-M_xtal_ dimer is rendered as surfaces with one protomer in green and the other in blue. (d) Superposition of the AlphaFold-predicted SARS2-M_AF_ monomer model (in orange) onto protomer A (in green) of batCOV5-M_xtal_ dimer. The batCOV5-M_xtal_ protomer B is displayed in blue. (e) Superposition of batCOV5-M_CTD-xtal_ (in orange) onto one protomer of batCOV5-M_xtal_ (in green). (f) Superposition of the cryo-EM structure of SARS2-ORF3a (in orange) onto batCOV5-M_xtal_ dimer (in green). For better viewing, each structure has one protomer rendered as cylinders.

The batCOV5-M_xtal_ structure is organized as a homodimer, with each protomer containing a short amino-terminus, a three-helix transmembrane domain (TMD) and a cytosolic CTD consisting of an eight-strand β-sandwich (Fig. 1b). The overall shape of the batCOV5-M_xtal_ dimer looks like a pair of pincers, with TMD as the “handle” and CTD as the “head” of the pincers (Fig. 1b). The two protomers form extensive TMD-TMD and CTD-CTD contacts, suggesting that the batCOV5-M_xtal_ dimer is a stable form (Fig. 1c). Not surprisingly, superposition of SARS2-M_AF_ onto one protomer of the batCOV5-M_xtal_ structure yielded an all-C_α_ RMSD of 4.57 Å with the major difference in TM1 (Fig. 1d). In the batCOV5-M_xtal_ structure, TM1 is placed almost in the sample plane as TM2/TM3, and the three helices of TMD resemble a capital letter “N”, while in SARS2-M_AF_ the TMD helices form a more compact three-helix bundle. As a result, the two TM1 helices swap positions in the batCOV5-M_xtal_ dimer compared to SARS2-M_AF_. Meanwhile, alignment between the batCOV5-M_CTD-xtal_ structure and CTD of the batCOV5-M_xtal_ structure yielded an all-atom RMSD of only 0.54 Å, indicating that the individually expressed CTD of batCOV5-M protein preserves its structure (Fig. 1e). This result may explain why molecular replacement for batCOV5-M_xtal_ failed when using SARS2-M_AF_ as a template, but succeeded with the batCOV5-M_CTD-xtal_ structure. Intriguingly, a structural comparison analysis using the Dali server^31^ indicates the batCOV5-M_xtal_ structure shares fold with the cryo-EM structure of SARS-COV-2 ORF3a protein (termed SARS2-ORF3a_EM_, PDB: 6XDC), which is a cation channel and also a homodimer^32^. Indeed, superposition of the two dimer structures yielded an all-C_α_ RMSD of 3.75 Å, suggesting that the overall folding of the two structures is alike (Fig. 1f). Nevertheless, few polar/charged residues in the batCOV5-M TMD (Supplementary Fig. 1) makes it less likely also a cation channel as ORF3a.

To validate the batCOV5-M_xtal_ structure and to study its function, we then investigated the interaction between M and N proteins (M-N interaction). The M-N interaction has been shown to facilitate virion assembly and budding of virus-like particles^6,7,13-15^. N protein is an RNA-binding protein containing two RNA-binding domains: an amino-terminal domain (N_1b_) and a carboxy-terminal domain (N_2b_)^13-15^ (Fig. 2a). An N_3_ region at the end of the carboxy-terminus has been shown to interact with M protein^13-15^. Therefore, we generated four constructs for expressing N protein and its fragments: the full-length protein (residues 1-427, termed batCOV5-N_FL_), the N_1_ fragment (residues 1-190, termed batCOV5-N_1_), the N_2_ fragment (residues 191-390, termed batCOV5-N_2_), and the N_3_ fragment (residues 391-427, termed batCOV5-N_3_) (Fig. 2a). In a pull-down analysis, purified batCOV5-N_FL_ and batCOV5-N_3_ proteins, but not batCOV5-N_1_ or batCOV5-N_2_, displayed robust interactions with purified wild-type batCOV5-M protein (Fig. 2b, lane 4 and 10), consistent with previous reports^13-15^. It has also been suggested that the N_3_ region may interact with M protein through electrostatic interactions^33-35^. Interestingly, batCOV5-N_3_ contains six basic residues in its amino-terminal half (residues 391-410, termed batCOV5-N_3N_) and five acidic residues in its carboxy-terminal half (residues 411-427, termed batCOV5-N_3C_) (Fig. 2a). The pull-down analysis clearly showed that purified batCOV5-N_3C_, but not batCOV5-N_3N_, pulled M protein down (Fig. 2b, lane 14). Furthermore, single mutations at the five acidic residue positions (E415, D416, D419, E424 and E426) of batCOV5-N_3_ decreased its binding affinity to batCOV5-M to varying extent in a microscale thermophoresis assay (Supplementary Table 1). This result is consistent with the notion that the negatively-charged batCOV5-N_3C_ interacts with the Lys/Arg/His-rich CTD of batCOV5-M (Supplementary Fig. 1), and charge interactions may play an important role in this process.

**Figure 2.**
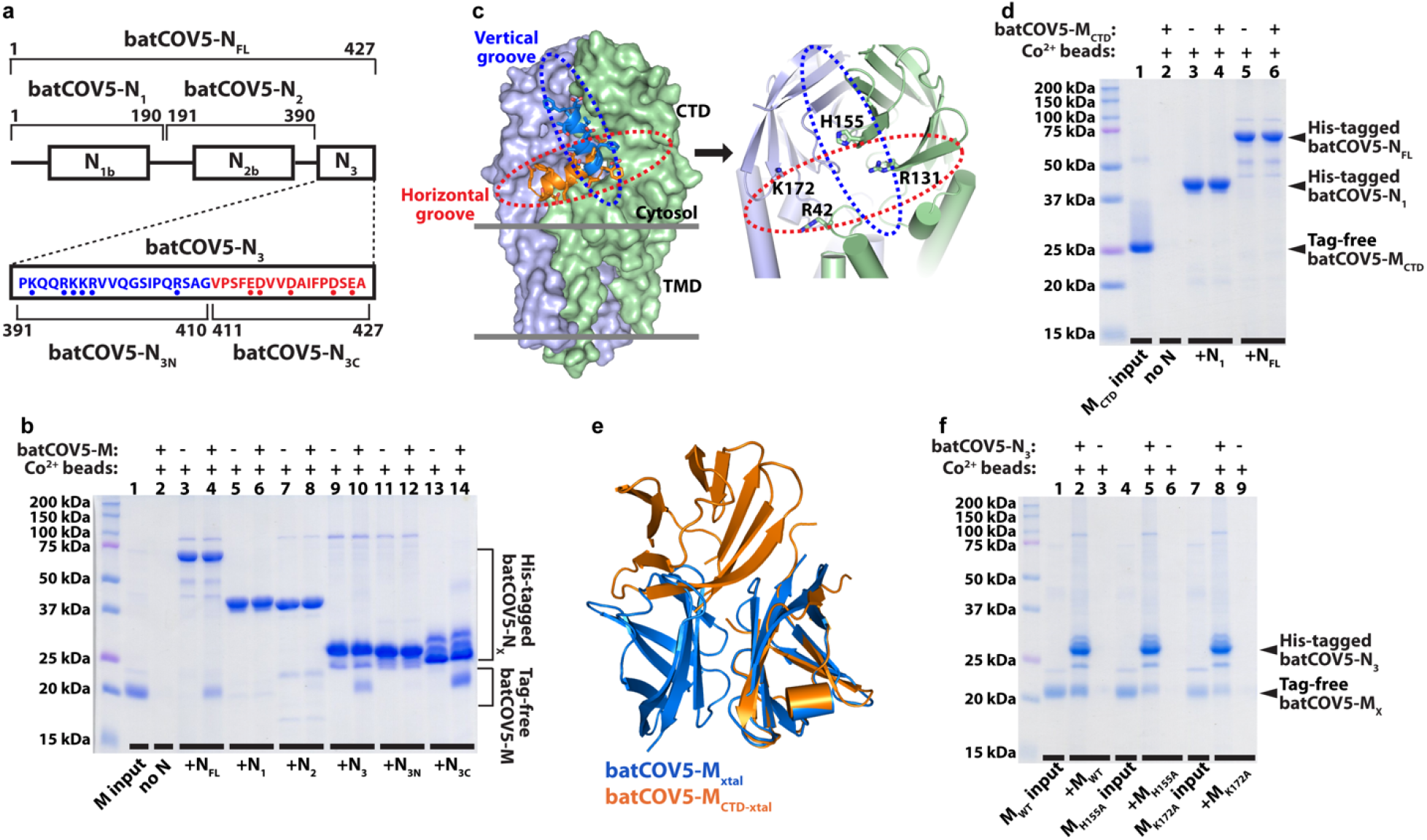
The interaction mechanism between M and N proteins of batCOV5. (a) Scheme of batCOV5-N fragments used in this study. The N_3N_ and N_3C_ sequences are shown with blue or red dots indicating basic or acidic residues, respectively. (b) A pull-down experiment using purified His-tagged batCOV5-N fragments to pull down tag-free batCOV5-M protein is examined by SDS-PAGE gel analysis. Signs and labels above and below the gel indicate the components present in the pull-down experiment. The label “batCOV5-N_X_” refers to all batCOV5-N fragments in this panel. Lane 1 is the batCOV5-M protein input for pull-down. Molecular weight of protein standards is labeled on the left of gel. (c) Left: Two docked models of batCOV5-N_3C_ (as an orange helix and a blue helix, respectively) into the batCOV5-M_xtal_ dimer structure. The two protomers are displayed in light blue and light green surfaces. The “horizontal groove” and the “vertical groove” are indicated by a red and a blue dashed oval, respectively. Right: An enlarged view of the “horizontal groove” and the “vertical groove”. Four basic residues within the two grooves are rendered as sticks. (d) A pull-down experiment using purified His-tagged batCOV5-N fragments to pull down tag-free batCOV5-M_CTD_-GFP protein, similar to panel b. To simplify text labels, “batCOV5-M_CTD_-GFP” is shortened as “batCOV5-M_CTD_” in this panel. (e) Superposition of batCOV5-M_CTD-xtal_ dimer (in orange) onto CTD of the batCOV5-M_xtal_ dimer (in blue). (f) A pull-down experiment using purified His-tagged batCOV5-N_3_ to pull down tag-free wild-type or mutant batCOV5-M proteins, similar to panel b. The label “batCOV5-M_X_” refers to all batCOV5-M variants in this panel. All pull-down experiments were repeated three times using biologically independent samples with similar results.

To further explore the binding mechanism of M and N_3C_ proteins, we created docking models using the batCOV5-M_xtal_ dimer structure and the batCOV5-N_3C_ sequence through the HDOCK web server^36,37^, which predicts batCOV5-N_3C_ as a helix. Two plausible binding patterns between M and N_3C_ were observed (Fig. 2c) after manually excluding incorrectly placed N_3C_ helix (e.g. in TMD). In the first pattern, the N_3C_ helix binds at the interface between the CTD dimer and the TMD dimer, which form a groove nearly horizontal to the membrane plane. In the second pattern, the N_3C_ helix binds to a groove formed along the CTD dimer interface nearly vertical to the membrane plane. Both binding patterns require dimerization of batCOV5-M. Consistently, individually-expressed batCOV5-M_CTD_ does not bind batCOV5-N_FL_ (Fig. 2d), likely due to that batCOV5-M_CTD_ does not form the same dimer interface as the full-length batCOV5-M does (Fig. 2e). Nonetheless, which binding pattern may be correct? To gain more insights, the basic residues within the two putative binding grooves of batCOV5-M were selected for mutagenesis.

For the “horizontal groove”, Arg42 and Lys172 were selected, whereas Arg131 and His155 were selected for both grooves as the two residues are located at the crossing of the two grooves (Fig. 2c, right panel). Although batCOV5-M mutants of R42A and R131A failed to express, the other two mutants (H155A and K172A) were expressed and showed apparently less binding to batCOV5-N_3_ in pull-down assays (Fig. 2f), which favors the “horizontal groove” for N_3C_ binding. Interestingly, His155 is not charged under the pull-down assay condition (pH 7.5), suggesting that interactions other than charge interactions likely also contribute to M-N binding. To verify the H155A result, binding of batCOV5-M_H155A_ to batCOV5-N_3_ was measured by microscale thermophoresis, which showed a decreased binding affinity (equilibrium dissociation constant *K*_d_ = 3.60 ± 0.17 µM) compared to the wild-type batCOV5-M protein (*K*_d_ = 0.77 ± 0.13 µM) (Supplementary Table 1), consistent with the pull-down result.

In this study, we report the first high-resolution crystal structure of M protein from a SARS-COV-2-related coronavirus batCOV5 and provide insight into the binding mechanism between M and N proteins. During preparation of this manuscript, Dolan and colleagues also reported a cryo-EM structure of M protein from SARS-COV-2 (termed SARS2-M_EM_) in bioRxiv^38^. As expected, SARS2-M_EM_ is also a homodimer and shares a similar fold to the batCOV5-M_xtal_ structure (Fig. 3a). Intriguingly, superposition of SARS2-M_EM_ dimer onto batCOV5-M_xtal_ dimer yielded a large all-C_α_ RMSD of 8.25 Å, suggesting that the two structures are in different conformations. Alignment of the two dimer structures showed an apparent difference in shape, with batCOV5-M_xtal_ dimer long and slim (∼80 Å x 45 Å) while SARS2-M_EM_ dimer short and fat (∼65 Å x 55 Å) (Fig. 3a). This observation is consistent with the “long” and “compact” conformations suggested previously^9^, which may be involved in M-N interaction and virus assembly. To further analyze the conformational change between SARS2-M_EM_ and batCOV5-M_xtal_, we superimposed single protomers of the two structures, yielding an all-C_α_ RMSD of 5.14 Å, whereas alignment of single CTDs and single TMDs between the two structures yielded all-C_α_ RMSDs of 1.38 Å and 4.18 Å, respectively (Fig. 3b). Comparison of single protomers between SARS2-M_EM_ and batCOV5-M_xtal_ reveals that their CTDs are highly alike, but their TMDs differ significantly in TM1, which is much closer to TM2 in SARS2-M_EM_ than in batCOV5-M_xtal_ (Fig. 3b). As a result, using TM2/TM3 as anchors, TM1 swings ∼40 degrees away from TM2 and CTD undergoes a rigid-body rotation of ∼30 degrees away from TMD during a transition from the “compact” SARS2-M_EM_ to the “long” batCOV5-M_xtal_ (Fig. 3b). In the meantime, the two protomers move ∼10 Å closer (Fig. 3a). In an analogy to the pincers, SARS2-M_EM_ dimer is like a pair of relaxed pincers, whereas batCOV5-M_xtal_ dimer is like the “handle” of the pincers is being clenched. It is also interesting to note that Trp24 of TM1 in batCOV5-M_xtal_ dimer form a π-cation-π interaction with an ammonium ion sitting between the two indole side groups (Fig. 3c), which may play a role in stabilizing the “long” conformation of M protein. Similarly, two tryptophan residues are also present in TM1 of SARS2-M (Trp20 and Trp31, Supplementary Fig. 1). It has been suggested that the “long” form, but not the “compact” form, of M protein binds N protein^9^. Therefore, it is desirable to determine an M-N complex structure in future studies to further elucidate the structure and function of M protein in coronaviruses.

**Figure 3.**
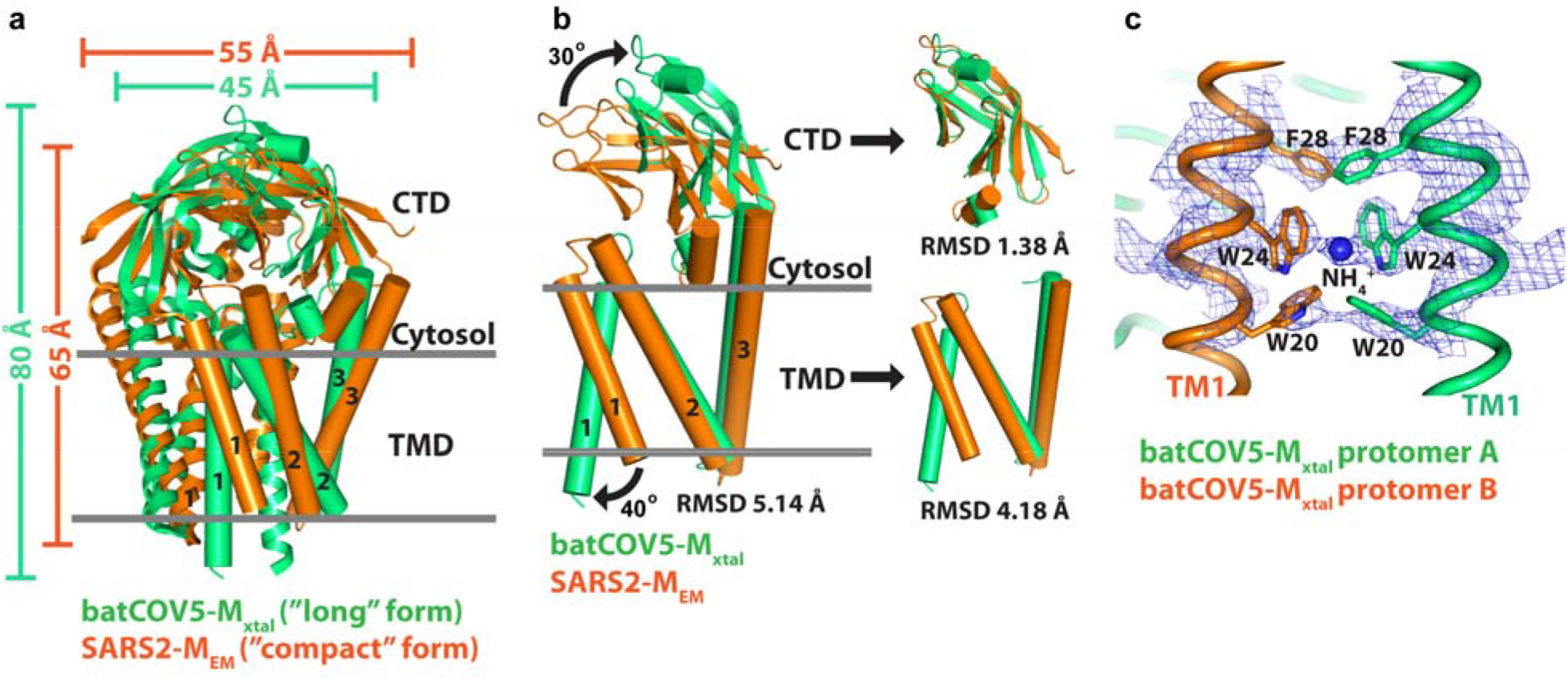
Structural comparison between batCOV5-M_xtal_ and SARS-M_EM_. (a) Superposition of the cryo-EM structure of SARS2-M (in orange) onto batCOV5-M_xtal_ dimer (in green). For better viewing, each structure has one protomer rendered as cylinders. Dimensions of the two dimers are indicated. (b) Left: Superposition of one SARS2-M_EM_ protomer (in orange) onto one batCOV5-M_xtal_ protomer (in green) using TM2/TM3 as anchors. Right: Individual CTDs and TMDs of the two structures are aligned using C_α_ atoms. (c) The simulated-annealing 2F_o_-F_c_ map contoured at 1.2 σ level is shown as blue mesh around an ammonium ion between the two Trp24 residues of TM1 in batCOV5-M_xtal_ dimer. The two protomers are displayed as tubes in orange and in green, respectively. The ammonium ion is rendered as a blue sphere and Trp24 as sticks.

## Online methods

### Protein expression and purification

The gene encoding M protein of *Pipistrellus* bat coronavirus HKU5 (batCOV5-M, NCBI accession YP_001039968.1) was synthesized (Genewiz, China) and cloned into a modified pPICZ plasmid (Thermo Fisher Scientific) containing an N-terminal tag of FLAG-His_10_-TEV protease recognition site. All batCOV5-M mutations were introduced by QuikChange II system (Agilent) according to manufacturer’s recommendation, and all mutations were verified by DNA sequencing. The constructs were linearized and transformed into *Pichia pastoris* strain GS115 by lithium chloride/single-strand carrier DNA/polyethylene glycol method according to manufacturer’s manual (Thermo Fisher Scientific). The transformants were inoculated into YPD medium consisting of 1% (w/v) yeast extract, 2% (w/v) peptone and 2% (w/v) D-(+)-glucose at 30 °C with shaking at 220 rpm until an OD_600_ of 3-5 was reached. To induce protein expression, yeast cells were harvested by centrifugation and resuspended to an OD_600_ of 1 in YPM medium consisting of 1% (w/v) yeast extract, 2% (w/v) peptone, 0.8% (v/v) methanol and 2.5% (v/v) dimethyl sulfoxide (DMSO) at 30 °C for 24 h. Cell pellets were resuspended in Lysis Solution (LS) containing 20 mM Tris-HCl pH 7.5, 150 mM NaCl, 10% (v/v) glycerol, 1 mM phenylmethanesulfonyl fluoride (PMSF) and 2 mM β-mercaptoethanol, and were lysed by an AH-1500 high-pressure homogenizer (ATS, China) at 1,300 MPa. Undisrupted cells and cell debris were separated by centrifugation at 3,000 x g, and membrane were collected by ultracentrifugation at 140,000 x g for 1 h at 4 °C. Protein was extracted by addition of 1% (w/v) n-dodecyl-β-D-maltopyranoside (DDM, Anatrace) at 4 °C for 2 h and the extraction mixture was centrifuged at 200,000 x g for 30 min at 4 °C. The supernatant was incubated with Co^2+^ resin in the presence of 20 mM imidazole pH 8.0 at 4 °C for 1 h, and the mixture was loaded in an empty chromatography column. The resin/protein was washed with 20 bed-volume of LS containing 2 mM DDM and 30 mM imidazole pH 8.0, and the protein was eluted with LS supplemented with 2 mM DDM and 250 mM imidazole pH 8.0.

To generate the batCOV5-M_CTD_-GFP construct, a superfolder GFP^29^ was split into two halves^30^ and was fused to the N- and C-termini of batCOV5-M_CTD_ (residues 115-203 of batCOV5-M) by gene synthesis (Genewiz, China). The fusion protein-encoding DNA was cloned into a modified pPICZ plasmid (Thermo Fisher Scientific) containing a C-terminal TEV site and a His_10_ tag. Transformation and expression of batCOV5-M_CTD_-GFP followed the same protocol as batCOV5-M except that the expression was induced at 25 °C. Cell pellets were resuspended in LS containing 20 mM Tris-HCl pH 8.0, 150 mM NaCl, 20% (v/v) glycerol and 1 mM PMSF, and were lysed similarly to batCOV5-M. Undisrupted cells and cell debris were separated by centrifugation at 140,000 x g at 4 °C for 1 h. The supernatant was supplemented with 20 mM imidazole pH 8.0 and was immediately loaded onto a pre-washed Co^2+^ affinity column. The column was then washed with 20 bed-volume of LS containing 30 mM imidazole pH 8.0, and the protein was eluted with LS containing 250 mM imidazole pH 8.0.

The gene encoding N protein of batCOV5 (batCOV5-N, NCBI accession YP_001039969.1) was synthesized (Genewiz, China) and cloned into a modified pPICZ plasmid (Thermo Fisher Scientific) containing an N-terminal tag of FLAG-His_10_-TEV site, followed by the bacterial cytochrome b562RIL (BRIL)^39^ to improve expression of the protein of interest. All batCOV5-N fragments were generated by polymerase chain reaction (PCR) and cloned into the same vector as batCOV5-N. For microscale thermophoresis (MST) analysis, batCOV5-N_3_-enconding DNA was sub-cloned into another modified pPICZ plasmid (Thermo Fisher Scientific) containing a C-terminal tag of TEV site-GFP-His_10_. Single mutations of batCOV5-N_3_ were introduced by QuikChange II system (Agilent). Transformation, expression and purification of batCOV5-N and fragments followed the same protocol as batCOV5-M_CTD_-GFP, except that LS contained 20 mM Tris-HCl pH 7.5, 150 mM NaCl, 10% (v/v) glycerol and 1 mM PMSF.

### Crystallization

Affinity-purified batCOV5-M protein was concentrated to 5 mg/ml, and treated with trypsin (TPCK-treated, Sigma-Aldrich) at a 1:50 ratio (trypsin:batCOV5-M, w/w) for 20 min at 18 °C to generate a stable core. The digestion was stopped by 10 mM PMSF and the protein was further purified by size-exclusion chromatography (SEC) in a buffer consisting of 150 mM NaCl, 20 mM Tris-HCl pH 7.5, 0.5 mM DDM and 5 mM β-mercaptoethanol. The peak fractions were pooled, concentrated to ∼30 mg/ml, and mixed with 1-oleoyl-rac-glycerol (monoolein, Sigma-Aldrich) at a 2:3 ratio (protein:lipid, w/w) using the twin-syringe mixing method^28^. The protein-lipid mixture was dispensed manually in ∼50-nl drops onto 96-well glass sandwich plates and overlaid with 0.8 μl of precipitant solution per drop. The batCOV5-M crystals were grown in 300 mM ammonium formate, 50 mM Tris-HCl pH 8.8, 35% (v/v) PEG 500 monomethyl ether and 7.1 mM pentaethylene glycol monooctyl ether. The crystals usually appear in one week and grow to full size in two weeks, and were flash-frozen directly in liquid nitrogen without additional cryoprotection.

For batCOV5-M_CTD_-GFP, affinity-purified protein was used directly for crystallization by vapor diffusion sitting-drop method without further SEC purification or concentration step. The batCOV5-M_CTD_-GFP crystals were grown in 0.4 M ammonium sulfate, 0.1 M Bis-Tris pH 5.3, 27% (w/v) PEG 3350 and 0.5% (v/v) ethyl acetate. The crystals were either cryoprotected by 15-20% (v/v) glycerol or directly flash-frozen in liquid nitrogen without additional cryoprotection.

### Data collection and structure solution

Diffraction data were collected on beamlines BL18U1 and BL19U1^40^ of National Facility for Protein Science in Shanghai (NFPS) at Shanghai Synchrotron Radiation Facility (SSRF). The data were indexed, integrated and scaled using the autoPROC pipeline package (Global Phasing Limited)^41^, which includes XDS^42^ and AIMLESS (CCP4 package)^43^. The batCOV5-M_CTD_-GFP structure was solved by molecular replacement with Phaser^44^ using a published superfolder GFP structure^29^ (PDB: 2B3Q) as a template. The full-length batCOV5-M structure was then solved by molecular replacement using the assembled batCOV5-M_CTD_ structure (Supplementary Fig. 2e) as a searching model. Manual model building and refinement was carried out using Coot^45^ and phenix.refine^46^, and Molprobity^47^ was used to monitor and improve protein geometry. Non-crystallographic symmetry (NCS) restraints were applied throughout the refinement to improve maps. The data collection and refinement statistics were generated using phenix.table_one^46^ and the values are listed in Supplementary Table 2. All structural figures, RMSD calculations and length/angle measurements were performed in PyMOL (Schr□dinger, LLC).

### Pull-down assay

For pull-down assays, affinity-purified batCOV5-M and mutants were treated with TEV protease (to remove His-tag) and Endoglycosidase H (New England Biolabs), and purified batCOV5-M_CTD_-GFP was treated with TEV protease, then all were further purified by SEC. In a pull-down experiment, affinity-purified batCOV5-N and fragments were first incubated with Co^2+^ beads for 30 min at 4 °C, followed by addition of His-tag-free batCOV5-M or variants to continue incubation for another 30 min at 4 °C in the presence of 30 mM imidazole pH 8.0. The Co^2+^ beads were collected by centrifugation at 5,000 x g for 1 min and washed twice with the assay buffer before being analyzed by SDS-PAGE gels.

### Docking of batCOV5-N_3C_ in batCOV5-M

Computational docking was performed using the HDOCK web server^36^ (http://hdock.phys.hust.edu.cn/). For receptor, the batCOV5-M structure was prepared as a .pdb file containing only one protein dimer (chain A and chain B) by removing other chains and all “HETATM” records. For ligand, non-polar residues at both ends of the batCOV5-N_3C_ sequence (VPSFEDVVDAIFPDSEA, 17 residues) were removed to generate a core sequence (SFEDVVDAIFPDSE, 14 residues), which was submitted directly to the system for template-free docking. Docked models were manually examined to remove incorrectly placed batCOV5-N_3C_ (e.g. in TMD).

### Microscale thermophoresis (MST)

MST analysis was performed using Monolith NT.115 (NanoTemper, Germany). All affinity-purified proteins were further purified by SEC in a buffer containing 20 mM Tris-HCl pH 7.5, 150 mM NaCl and 0.5 mM DDM. The peak fractions were pooled and diluted to 100 nM using the SEC buffer. The GFP moiety fused to batCOV5-N_3_ and its mutants provided the fluorescence signal required by MST. Meanwhile, purified batCOV5-M and batCOV5-M_H155A_ served as ligands and were prepared according to the MST manual with the highest concentration of 32 μM for both proteins. The batCOV5-N_3_ and mutant samples were mixed with serial-diluted ligands and were incubated for 10 min at room temperature. Then the samples were loaded into capillaries and MST measurements were performed according to the Monolith manual. The equilibrium dissociation constant (*K*_d_) was determined in the MO.Affinity Analysis software (NanoTemper, Germany) with the *K*_d_ fit function. All MST measurements were performed in three biologically independent experiments (*N*=3), and *K*_d_ values are expressed as mean ± SD in the text and Supplementary Table 1. Two-tailed Student’s t-test was performed for statistical analysis in Supplementary Table 1.

## Acknowledgements

Diffraction data used in this study were collected on beamlines BL18U1 and BL19U1 of National Facility for Protein Science in Shanghai (NFPS) at Shanghai Synchrotron Radiation Facility (SSRF). The authors thank the staff from these beamlines for assistance during data collection. This work was supported in part by the National Natural Science Foundation of China (NSFC) grant 31770783, and the 1.3.5 Project for Disciplines of Excellence grant by West China Hospital of Sichuan University to XZ.

## Data availability

The atomic coordinates and structure factors generated in this study have been deposited in the Protein Data Bank under the following accession codes: 7Y96 for batCOV5-M_CTD_-GFP and 7Y9B for batCOV5-M. Three previously reported structures used in this study are available in the Protein Data Bank under the following accession codes: 2B3Q for a superfolder GFP, 6XDC for ORF3a of SARS-COV-2 and 8CTK for M protein of SARS-COV-2. Source data of *K*_d_ values are provided in Supplementary Table 1.

## Author contributions

XZ and ZS conceived and supervised the project, and wrote the manuscript with input from all authors. XW performed structural and functional studies; YY performed functional studies; and all authors analyzed the data.

