## Supplementary information for "Crystal structure of the membrane (M) protein from a SARS-COV-2-related coronavirus"

### Supplementary tables

#### Supplementary Table 1. Affinities of batCOV5-M to batCOV5-$N_3$ determined by MST.

| ($K_d$: μM) | Repeat 1 | Repeat 2 | Repeat 3 | Mean | SD | *P* |
|---|---|---|---|---|---|---|
| **M WT/$N_3$ WT** | 0.65 | 0.77 | 0.90 | 0.77 | 0.13 | - |
| **M WT/$N_3$ E415A** | 3.40 | 3.40 | 3.70 | 3.50 | 0.17 | < 0.001 |
| **M WT/$N_3$ D416A** | 3.00 | 3.00 | 3.20 | 3.07 | 0.12 | < 0.001 |
| **M WT/$N_3$ D416N** | 2.10 | 2.20 | 2.70 | 2.33 | 0.32 | 0.001 |
| **M WT/$N_3$ D419A** | 4.50 | 5.00 | 5.00 | 4.83 | 0.29 | < 0.001 |
| **M WT/$N_3$ E424A** | 1.20 | 1.90 | 2.60 | 1.90 | 0.70 | 0.052 |
| **M WT/$N_3$ E426Q** | 2.80 | 3.40 | 3.80 | 3.33 | 0.50 | 0.001 |
| **M H155A/$N_3$ WT** | 3.40 | 3.70 | 3.70 | 3.60 | 0.17 | < 0.001 |

All MST measurements were repeated with three biologically independent samples (*N*=3) and source data of $K_d$ values are shown. Two-tailed Student's t-test was performed between the "M WT/$N_3$ WT" group and other groups, and *P* values are shown.

**Supplementary Table 2. Data collection and refinement statistics.**

| | batCOV5-$M_{CTD}$-GFP | batCOV5-$M_{xtal}$ |
|---|---|---|
| PDB ID | 7Y96 | 7Y9B |
| **Data collection** | | |
| Space group | P $6_1$ 2 2 | P 1 $2_1$ 1 |
| Wavelength (Å) | 0.97915 | 0.97915 |
| Unit cell | | |
| a, b, c (Å) | 82.02, 82.02, 427.84 | 76.57, 66.57, 112.63 |
| $\alpha$, $\beta$, $\gamma$ (°) | 90, 90, 120 | 90, 109.83, 90 |
| Resolution (Å) | 3.42 (3.54-3.42) | 3.21 (3.33-3.21) |
| Unique reflections | 11763 (1138) | 16670 (1745) |
| Multiplicity | 4.4 (4.6) | 3.4 (3.4) |
| Completeness (%) | 93.77 (96.11) | 93.93 (98.75) |
| $I/\sigma I$ | 9.63 (3.09) | 7.42 (1.76) |
| $R_{merge}$ | 0.147 (0.508) | 0.141 (0.694) |
| $R_{meas}$ | 0.166 (0.569) | 0.167 (0.827) |
| $R_{pim}$ | 0.075 (0.252) | 0.090 (0.444) |
| $CC_{1/2}$ | 0.99 (0.841) | 0.994 (0.74) |
| **Refinement** | | |
| Resolution (Å) | 3.42 (3.76-3.42) | 3.21 (3.46-3.21) |
| No. reflections | 11762 | 16586 |
| Completeness (%) | 93.9 | 94.0 |
| $R_{work}/R_{free}$ (%) | 24.0/26.5 | 27.2/29.6 |
| No. atoms | 4890 | 6041 |
| Protein | 4846 | 5939 |
| Ligands | 44 | 98 |
| Solvent | | 4 |
| Average B-factor | 65.08 | 58.37 |
| Protein | 65.19 | 58.48 |
| Ligands | 53.02 | 51.76 |
| Solvent | | 56.47 |
| Ramachandran | | |
| Favored (%) | 97.54 | 98.38 |
| Allowed (%) | 2.46 | 1.62 |
| Outliers (%) | 0.00 | 0.00 |
| RMS bonds (Å) | 0.003 | 0.002 |
| RMS angles (°) | 0.658 | 0.425 |
| Clashscore | 8.73 | 6.70 |

Statistics for the highest-resolution shell are shown in parentheses.

### Supplementary figures

```
                                     TM1
MERS-M        M-SNMTQLTEAQIAIIKDWNFAWSLIFLLITIVLQYGYPSRSMTVYVFKMFVLWLLWPS    59
batCOV5-M     MASSNVTLSNDEVLRLVKDWNFTWSVVFLLITIVLQYGYPSRSMFVYVIKMFVLWLLWPA   60
SARS2-M       MADSNGTITVEELKKLLEQWNLVIGFLFLTWICLLQFAYANRNRFLYIIKLIFLWLLWPV   60
SARS-M        MAD-NGTITVEKLKQLLEQWNLVIGFLFLAWIMLLQFAYSNRNRFLYIIKLVFLWLLWPV   59
              * .    ::  ::  ::::**:. ..:**    :**:.*..*.  :*::*:..******

                  TM2                TM3
MERS-M        SMALSIFSAIYPIDLASQIISGIVAAVSAMMWISYFVQSIRLFMRTGSWWSFNPETNCLL  119
batCOV5-M     SMALSIFCAVYPIDLASQIISGILAATSCAMWISYFVQSIRLFMRTGSWWSFNPESNCLL  120
SARS2-M       TLACFVLAAVYRINWITGGIAIAMACLVGLMWLSYFIASFRLFARTRSMWSFNPETNILL  120
SARS-M        TLACFVLAAVYRINWVTGGIAIAMACIVGLMWLSYFVASFRLFARTRSMWSFNPETNILL  119
              ::*  ::.*:* *:  :  *:  :*.    **:***: *:*** ** * ******:* **

MERS-M        NVPIGGTTVVRPLVEDSTSVTAVVTNGHLKMAGMHFGACDYDRLPNEVTVAKPNVLIALK  179
batCOV5-M     NVPIGGTTVVRPLVEDSTSVTAVVTDGYLKMAGMHFGACDFQRLPSEVTVAKPNVLIALK  180
SARS2-M       NVPLHGTILTRPLLESELVIGAVILRGHLRIAGHHLGRCDIKDLPKEITVATSRTLSYYK  180
SARS-M        NVPLRGTIVTRPLMESELVIGAVIIRGHLRMAGHSLGRCDIKDLPKEITVATSRTLSYYK  179
              ***: ** :.***:*..  : **:  *:*::**  :* ** . **.*:***....*   *

MERS-M        MVKRQSYGTNSGVAIYHRYKAGNYRS--PPITADIELALLRA                    219
batCOV5-M     MIKRQAYGTNSGVAIYHRYKAGNYRR--PPIIQDQELALLRA                    220
SARS2-M       LGASQRVAGDSGFAAYSRYRIGNYKLNTDHSSSSDNIALLVQ                    222
SARS-M        LGASQRVGTDSGFAAYNRYRIGNYKLNTDHAGSNDNIALLVQ                    221
              :   *  . :**.* * **: ***:        . ::***
```

**Supplementary Figure 1. Sequence alignment of several betacoronavirus M proteins by ClustalW[1,2].** TM helices are indicated by cylinders and labeled from TM1 to TM3. The batCOV5-$M_{CTD}$ sequence is highlighted in cyan. Tryptophan residues within TM1 are highlighted in green. Basic residues selected from the "horizontal groove" and the "vertical groove" in this study, including R42, R131, H155 and K172, are highlighted in magenta. Asterisks (*) indicate identical residues. Colons (:) indicate strong similarities. Periods (.) indicate weak similarities. MERS-M, MERS-COV M protein; batCOV5-M, *Pipistrellus* bat coronavirus HKU5 M protein; SARS2-M, SARS-COV-2 M protein; SARS-M, SARS-COV M protein.

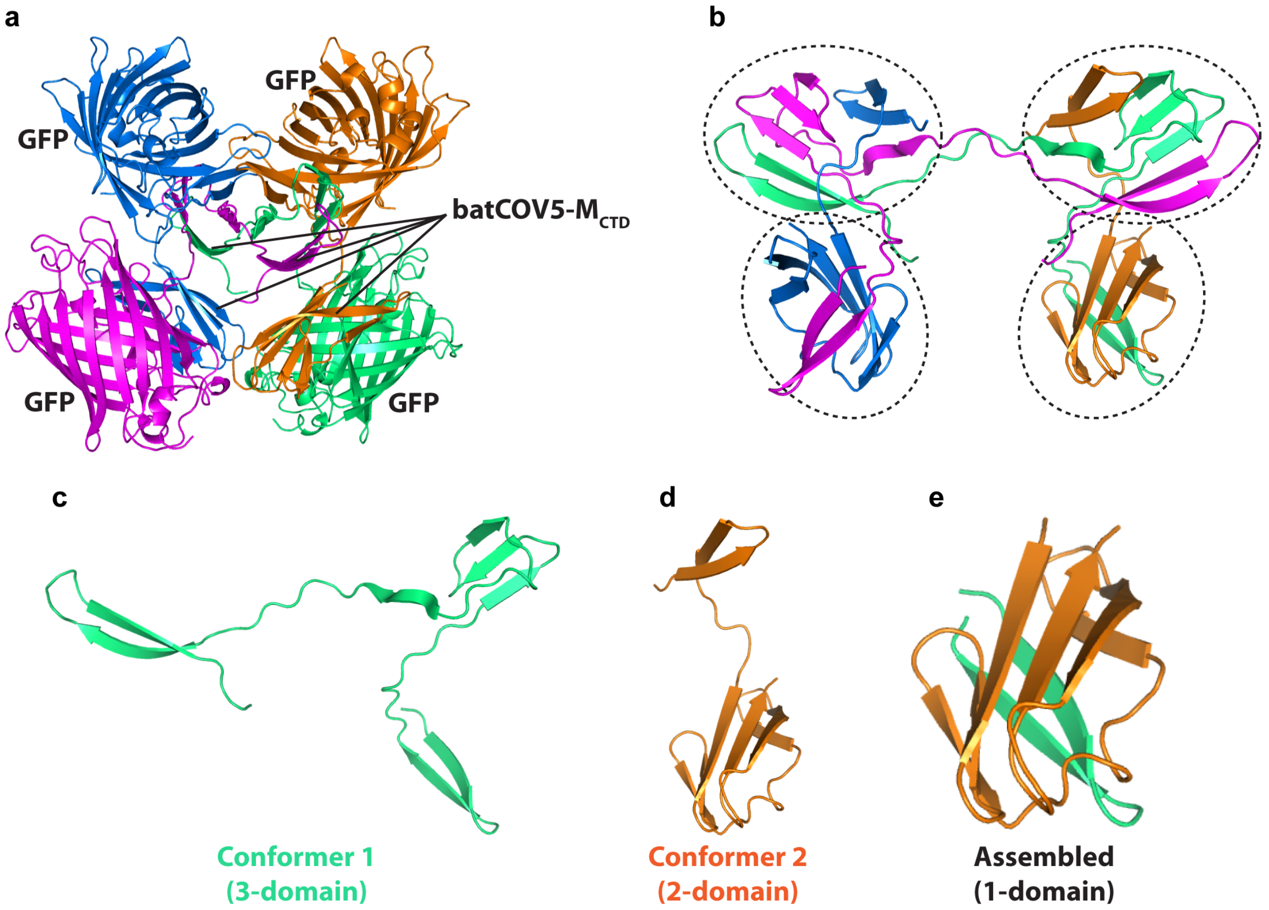


**Supplementary Figure 2. Crystal structure of batCOV5-M$_{CTD}$ fused with a superfolder GFP.** (a) Tetrameric batCOV5-M$_{CTD}$-GFP displayed in four colors. (b) The batCOV5-M$_{CTD}$ tetramer formed by swapping β strands with each other. Each color indicates one protomer. Each dashed oval indicates one assembled batCOV5-M$_{CTD}$. GFP is not displayed for clearer viewing. (c)-(e) Three conformations of batCOV5-M$_{CTD}$.
